# Translocating *Mycobacterium ulcerans*: an experimental model

**DOI:** 10.1101/2020.03.04.977447

**Authors:** N Hammoudi, M Fellag, M Militello, A Bouam, M Drancourt

## Abstract

*Mycobacterium ulcerans* is a non-tuberculous environmental mycobacterium responsible for extensive cutaneous and subcutaneous ulcers in mammals, named Buruli ulcer in patients. *M. ulcerans* has been seldom detected in the feces of mammals but not in patients, nevertheless the detection and isolation of *M. ulcerans* in animal feces does not feet with the current epidemiological schemes for the disease. Here using an experimental model in which rats were fed with 10^9^ colony-forming units of *M. ulceran*s, we detected *M. ulcerans* in feces of challenged rats for two weeks and along their digestive tract for 10 days. *M. ulcerans* was further detected in the lymphatic system including cervical and axillary lymph nodes and the spleen, but not in any other tissue including the healthy and breached skin, 10 days post-challenge. These observations indicate that in some herbivorous mammals, *M. ulcerans* contamination by the digestive route may precede translocation and limited infection of the lymphatic tissues without systemic infection. These herbivorous mammals may be sources of *M. ulcerans* for exposed populations but are unlikely reservoirs for the pathogen.

## INTRODUCTION

*Mycobacterium ulcerans* is an environmental slowly growing, non-tuberculous mycobacterium responsible for progressively extending cutaneous and subcutaneous ulcer named Buruli ulcer [1]. Buruli ulcer is a World Health Organization-notifiable neglected infection which has been notified by 34 countries over the last ten years [2]. Accordingly, Buruli ulcer is a tropical infection mainly affecting rural populations in South America, West Africa, Australia, South-China and Japan [3]. Although a genetic trait has recently been described among a 7-individual family in whom two individuals suffered Buruli ulcer and carried a specific deletion on chromosome 8 [4], in addition to previously reported deletion in the NRAMP-1 gene [5], nevertheless Buruli ulcer is not a contagious infection but is rather resulting from contacts with *M. ulcerans*-contaminated environments [2]. Accordingly, it’s been reported that *M. ulcerans* is being cultivated from aquatic Hemiptera [6]. indeed *M. ulcerans* DNA has also been detected in some animals including *Thryonomys swinderianus* (here designed agoutis, the word commonly used in West Africa) [7–8] rabbits and rats, which have all in common to be rodent mammals [9]. Moreover, two isolates of *M. ulcerans* have been reported from possum feces collected in Melbourne, region, Australia although characterization and repository reference were not provided [10]. These observations led to suggest that possums and *Pseudocheirus peregrinus* may play a role in the epidemiology of Buruli ulcer in endemic Australian regions [11].

Furthermore, we recently reported the isolation of *M. ulcerans* from *T. swinderianus* feces collected in the vicinity of the Kossou Dam, Côte d’Ivoire, although we were unable to sub-culture this isolate [8]. These observations suggested that *T. swinderianus*, an herbivorous mammal rodent may contaminate its digestive tract by the oral route after eating *M. ulcerans*-contaminated food; possibly acting as a secondary source of infection for populations as it is catched as brush meat and eaten after unprotected, manual evisceration [7] Likewise, it has been suggested that the small terrestrial mammals *Mastomys natalensis* can play a potential role in the natural history of *M. ulcerans* [12]. Also, we recently observed the PCR-based detection of the ketoreductase B gene (KR-b) and the IS*2404* and IS*2606* insertion sequences in one tenth of spleen specimens collected from *T. swinderianus* in the area of Yamoussoukro area, Côte d’Ivoire, and in spleen of common ringtail possums in some areas of Victoria endemic for *M. ulcerans* disease [7–10].

Altogether, these observations pointed to the medical interest in understanding the mode of contamination and pathology of *M. ulcerans* in *T. swinderianus*. We therefore developed an experimental model of oral route contamination in rat, asking four questions relative to the survival and excretion of the pathogen in the digestive tract, its translocation, its systemic dissemination and its capacity to infect previous aseptic skin lesion.

## MATERIALS AND METHODS

### Ethics Statements

The experimental protocol, registered by the “Ministère de l’Enseignement Supérieur et de la Recherche” under reference number 2018081011226001, was approved by the Institutional Animal Care and Use Committee of Aix-Marseille University “C2EA-14”, France. All animal handling was carried out in compliance with the rules of Décret N° 2013–118, Février 7, 2013, France. The experimental procedures on rats were carried out in accordance with European law and in agreement with Animal: Reporting In Vivo Experiments (ARRIVE Guidelines http://www.nc3rs.org.uk). We used Long-Evans rats (Charles River Laboratories, L’Arbresle, Lyon, France). Animals were housed in protected environmental area, in individual transparent cages (one rat per cage) in individually ventilated Allentown technologies caging systems (Allentown, Pennsylvania, USA) with free access to standard diet including dehydrated rodent feed pellets and sterile water until the experiment. Efforts have been made to minimize the number of animals, and to limit their stress the environment has been enriched with litter and cardboard tunnels. The rats were observed daily for any signs of distress or adverse events. To minimize animals suffering all invasive procedures have been performed under full general anesthesia. The animals were sacrificed by injection of lethal dose of Pentobarbital preceded by full general anesthesia. All experiments were performed in a biosafety level 3 laboratory of the University Hospital Institute (IHU), Marseille, France.

### *M. ulcerans* inoculum

*M. ulcerans* strain CU 001, a clinical isolate from Ghana [13] was cultured on Middlebrook 7H10 supplemented by OADC during a 6-week incubation at 30°C. Colonies were suspended on sterile phosphate buffered saline (PBS) tube and the bacterial was vigorously vortexed for 10 min using 3-mm sterile glass beads (Sigma-Aldrich, Saint-Quentin-Fallavier, France) and passed three times through a 29 Gauges in order to eliminate bacterial aggregates. The mycobacterial suspension was then calibrated at optic density of 5 McFarland (equivalent to 10^9^ colony-forming units (CFU)/mL.

### Rat infection protocol

Animal experimentations were performed on 16 rats (8 males and 8 females) aged 8 weeks (Charles River Laboratories) weighing between 220 g and 250 g. These animals obtained with complete health reports, were found to be healthy and free of infection and were housed under conditions free of specific pathogens. Each rat was placed into an individual plastic cage with free access to water and food. In order to induce subcutaneous lesions in the shoulders, rats were full general anaesthetized with intraperitoneal injection of a mixture of Ketamine (80 mg/kg) and Xylazine (10 mg/kg). One gram of sterile glass powder mixed with 300 μL of sterile PBS were injected under the skin of the right shoulder and 300 μL of PBS under the skin of the left shoulder as a negative control. Digestive inoculation was performed after the shoulder skin lesions resolved (i.e., 7 days after skin lesions were made). Briefly, the rats were manually restrained and sterile, single-use 1-mL syringes were used to administer the *M. ulcerans* suspensions directly into the rats’ mouth, respecting the swallowing cycle in order to ensure that rats swallowed the entire administered suspension: 300 μL of sterile PBS were administered to 4 rats (2 males and 2 females) forming the negative control group and 300 μL of mycobacterial suspension at 10^9^ CFU/mL were administered to 12 challenged rats (6 females and 6 males).

### Animal follow-up and sample collection

Rat behavior was observed daily until the day of euthanasia. In the post-infectious period, animals were observed carefully daily for any abnormal behavior including swelling/bleeding of the injection site, ruffled coat, hunched posture, signs of pain or distress. Feces were collected directly from the exit of the rectum on the first day of challenge and then every two days until 20 days after challenge. Ten days after challenge, rats of group n°1 of 6 infected rats (3 males and 3 females) and two controls (1 male and 1 female) were sacrificed by intraperitoneal injection of lethal dose of Pentobarbital (100-150 mg/kg) preceded by full general anesthesia as previously described. Organs were carefully collected.

Another group n°2 of 6 infected rats (3 males and 3 females) and one male and one female control rats were followed in the same way until 60 days after infection, then sacrificed by intraperitoneal injection of a mixture intraperitoneal injection of lethal dose of Pentobarbital (100-150 mg/kg) preceded by full general anesthesia as previously described above and euthanized to collect a second set of organs.

### PCR detection of *M. ulcerans* DNA

All collected organs were stored in Eppendorf tubes at 4°C for one day. Then, one piece of each organ was transferred to another 1.5 mL-Eppendorf tube containing 500 μL of sterile PBS. The organs were vigorously crushed using single used sterile piston and 200 μL of organ juice were put into a 1.5 mL-Eppendorf tube containing a mixture of 200 μL G2 lysis buffer, 20 μL proteinase K and a small quantity of glass powder. Tube underwent 3 cycles of FastPrep 24^TM^-5 (MP Biomedicals, Strasbourg, France) before being heated at 56°C for 2 hours. Then, 200 μL of supernatant were used to extract DNA using the EZ1 apparatus according to the manufacturer’s recommendations (Qiagen, GmbH, Germany). Extracted DNA was stored at 4°C. Detection of *M. ulcerans* DNA was performed by using real-time PCR (RT-PCR) and a CFX thermal cycler (BIO-Rad, Marnes-la-Coquette, France) using specific primers targeting the ketoreductase B gene (KR-b) and the IS*2404* and IS*2606* insertion sequences, as previously described [14]. The negative controls of our RT-PCR reactions were formed from the same reaction mix as our samples switching only the 5 uL of DNA by 5 uL of ultrapure™ DNase/RNase-Free Distilled Water (Invitrogen France), thus one negative control is placed after every 5 samples on a Light cycler 480 multiwell plates 96-well plate (Roche).

### *M. ulcerans* culture

One piece of spleen collected from every rat, was crushed in 500 μL of sterile PBS using sterile single-used piston and 2 × 150 μL of spleen juice were inoculated on two Middlebrook 7H10 agar plates incubated at 30°C for 2 months. In addition, the feces collected at 3, 14- and 20-days post-infection were incubated for 10 min at room temperature in 2 mL NAOH (1M) and centrifuged at 3,500 g for 10 min. The pellet was resuspended in 1 mL of 5% oxalic acid and incubated for 10 min at room temperature before centrifugation at 3,500 g for 10 min. The pellet was then resuspended into 2 mL home-made Trans MUl decontamination-preculture medium incubated for 5 days at 30°C. The mixture was then vortexed 30 times and 200 μL were plated onto Trans MUg, a homemade Middlebrook-based medium supplemented with with 10% oleic acid, bovine albumin, dextrose and catalase enrichment (OADC, Becton Dickinson), oxytetracycline (40 μg/mL), polymyxin E (80 μg/mL) and voriconazole (50 μg/mL) for 3 months at 30°C. Trans MUg medium was previously tested and shown to respect the viability of *M. ulcerans* (no published data).

## RESULTS

### Rat clinical follow-up

During a 60-day follow-up post-infection, all rats in the control group and all rats in the challenged group remained apparently healthy and showed no pathological clinical sign, no pain and no weight loss. In addition, the subcutaneous lesions induced by the glass powder do not show any pathological signs during the duration of the experimentation.

### RT-PCR test results

In all RT-PCR assays, the negative controls remained negative. At 10 days post-challenge, the RT-PCR detection of *M. ulcerans* DNA remained negative on the internal organs and feces collected in control group rats whereas it was positive in some *M. ulcerans*-infected rats: RT-PCR tests revealed a simultaneous detection of IS*2606* insertion sequence and the KR-B gene in cervical lymph nodes, axillary lymph nodes, spleen and the digestive tract in males and female (Table 1). Also, *M. ulcerans* DNA was detected in the feces of infected rats up to 15- and 17-days post-infection, in males and females, respectively. At 60 days post-challenge, the RT-PCR detection of *M. ulcerans* DNA remained negative on the internal organs and in feces collected in control and challenged groups (Table 1).

**Table 1.**
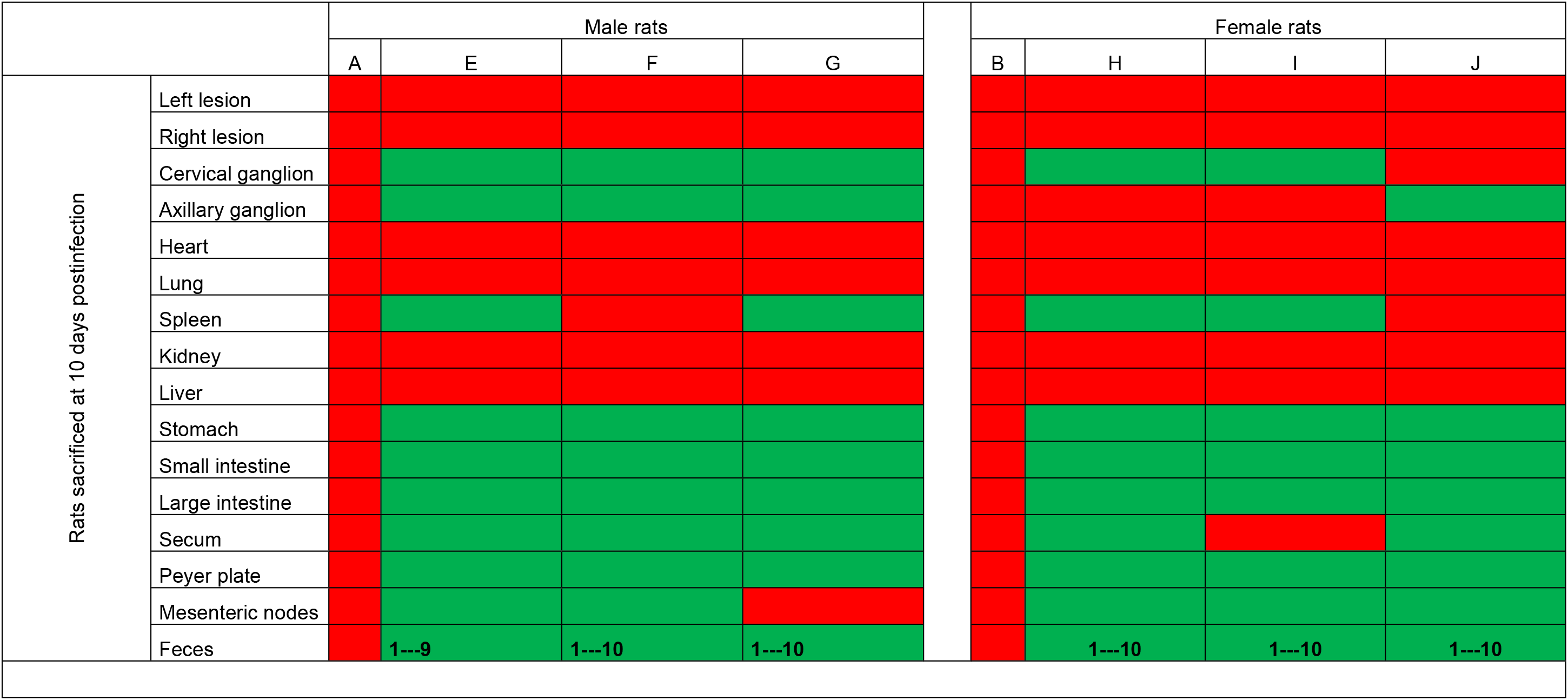
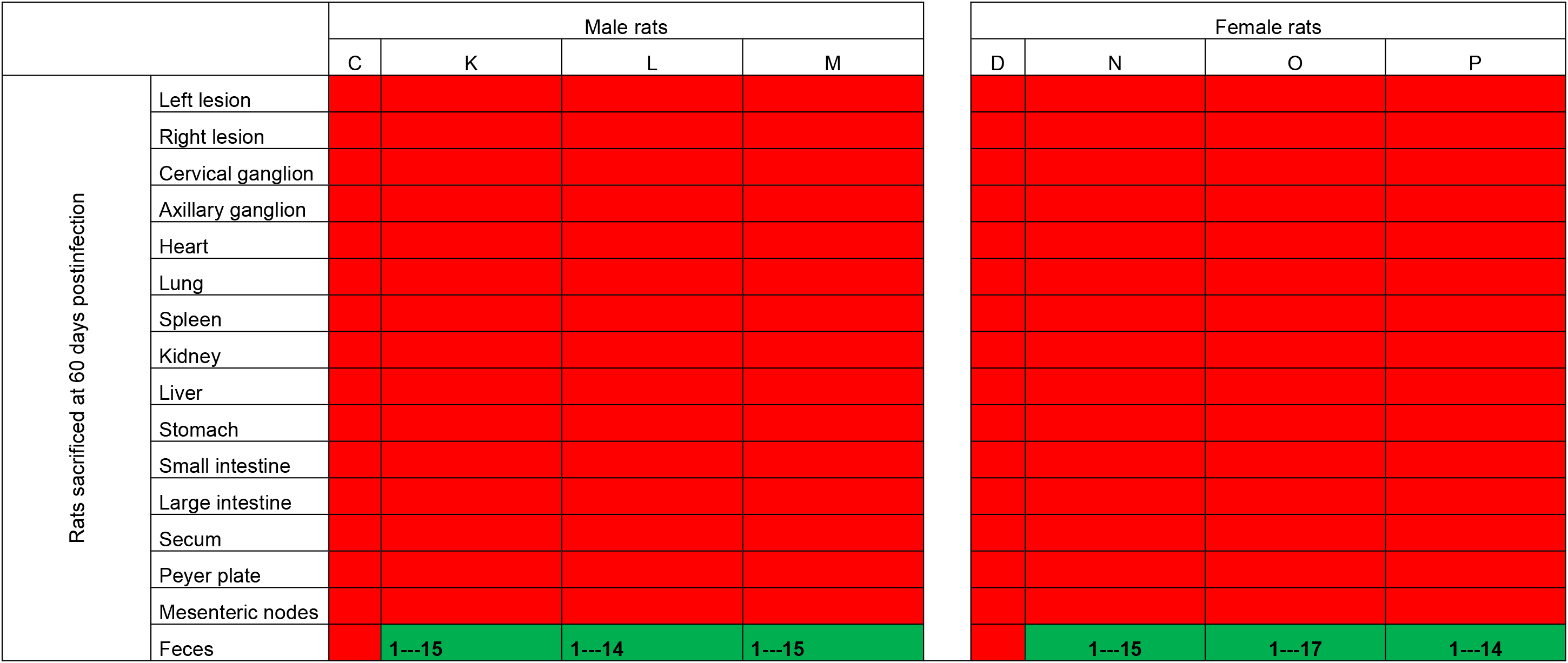
Detection of *M. ulcerans* in the organs and feces collected in rats challenged with the pathogen by the oral route. Red squares indicate the absence of detection, green squares indicate a positive detection. “A-B-C-D” denote negative control, non-challenged rats; “E-P” denote challenged rats. The numbers indicate the time (days) when M. ulcerans DNA was detected in the feces.

### Culture results

No colony of *M. ulcerans* was isolated from the feces and the spleen samples collected in 4 negative-control group rats after three-month incubation. The same observations were reported after three-months of cultivation of feces and spleens of 12 *M. ulcerans*-infected rats

## DISCUSSION

We have previously reported the molecular detection of *M. ulcerans* DNA in the digestive tract and the spleens of wild agoutis caught in Côte d’Ivoire: and its culture in one case [7–8] In the present work, we developed an experimental model of gastrointestinal infection in rats to confirm that our field observations were the result of probable gastrointestinal contamination of wild agoutis. We chose the rat model as the laboratory mammal closest to agouti, sharing its general morphology (in terms of size and weight), a herbivorous rodent diet and a body temperature of 37°C, which also makes it a relevant model of human infection [15].

The rat model showed that these rodents can be infected by *M. ulcerans* through the digestive tract, allowing us to answer our first question. Indeed, the negative control rats and the experimental negative controls that we introduced at all stages of our experiments, remained negative in both the RT-PCR and the culture-based experiments carried out on the feces and internal organs RT-PCR assays of uninfected rats. This fact allowed us to interpret the positive results we observed, as authentic and not a mere result of laboratory contamination.

Our experimental results thus confirm that some wild rodents are digestively infectable with *M. ulcerans,* as it has been previously observed for wild agoutis in Côte d’Ivoire [7–8]. and for possums in Australia [10]. Accordingly, our results reporting the *M. ulcerans* DNA detected in feces and in internal organs, suggests that some wild rodents may participate in the natural cycle of the pathogen. This observation indicates that direct contact with the feces of these wild rodents, when they are prepared as bushmeat for example, constitutes a circumstance of contamination of populations by live *M. ulcerans*, and a source of Buruli ulcer.

In a second step, *M. ulcerans* was detected by RT-PCR but not cultured, in the spleen and some lymph nodes of challenged rats but not from negative control rats. This observation indicates that *M. ulcerans* has probably the ability to penetrate the digestive mucosa through mechanisms whose determination was not part of the objectives of this experimental work, and to be kept by the lymphatic system in which it is destroyed. Accordingly, this observation suggests that at some stage of this translocation process, mycolactones are no longer synthesized, or secreted or inactivated. This experimental result corroborates the fact that *M. ulcerans* has never been detected in a tissue or organ at a distance from its inoculation point, which also suggests that the cytotoxic activity of mycolactones is local and not systemic, without clarifying the role of mycobacterial inhibition and mycolactone inhibition themselves. Interestingly, it is possible to determine mycolactones in Buruli ulcer lesions, but not in patients’ blood [16]. It should be noted that translocation property is shared with mycobacteria of the *Mycobacterium tuberculosis* complex, in which experimental translocation has been shown for *Mycobacterium canettii* [17] and *M. tuberculosis* [18], there are no experimental data or clinical observations to our knowledge for mycobacteria of the *Mycobacterium leprae* complex.

After translocation, the fate of *M. ulcerans* differs considerably from that of mycobacteria of the *M. tuberculosis* complex. The latter spread into the lung and other highly vascularized organs [18], while *M. ulcerans* did not spread into any organs in the rat model.

Altogether, the observations here reported indicate that in some herbivorous mammals, *M. ulcerans* contamination by the digestive route may precede translocation and limited infection of the lymphatic tissues without systemic infection. These herbivorous mammals may participate as sources of *M. ulcerans* for exposed populations, but do not participate as reservoirs for the pathogen.

## ACKNOWLEDGMENTS

NH obtained a PhD grant from the Fondation Méditerranée Infection, Marseille, France. This work was supported by Agence Nationale de la Recherche (ANR-17-CE35-0006-01 PRIME, http://www.agencenationale-recherche.fr/) and by the French Government under the « Investissements d’Avenir » (Investments for the Future) program managed by the ANR (reference: Méditerranée Infection 10-IAHU-03), as well as the Agence Nationale de la Recherche (ANR-17-CE35-0006-01 PRIME, http://www.agencenationale-recherche.fr/). This work was supported by Région Provence Alpes Côte d’Azur and European funding FEDER IHUBIOTK.

